# Evidence for an efferent-based prediction contributing to implicit motor adaptation

**DOI:** 10.1101/2024.07.29.605662

**Authors:** Annika Szarka, Hyosub E. Kim, J. Timothy Inglis, Romeo Chua

## Abstract

Models of sensorimotor adaptation have proposed that implicit adaptation is driven by error signals created by discrepancies between various sensory information sources. While proprioception has been suggested as a critical source for the error signals driving adaptation, the role of an efferent-based motor prediction has largely been neglected. In this study, we examined the effect of dissociating the afferent and efferent information available during implicit adaptation. Participants moved a visual cursor towards targets by applying horizontal forces to a stationary handle at a central home location. During perturbation trials, the cursor followed an invariant path rotated relative to the target. Participants were instructed to ignore this task-irrelevant cursor feedback and to isometrically “reach” towards the target. Participants implicitly adapted in the isometric task, even when the hand never actually moved to the target. Moreover, the level of adaptation surpassed that of a typical clamped reaching paradigm by nearly twofold. This was confirmed in a secondary experiment where participants performed actual reaching movements and demonstrated significantly less adaptation. Our findings suggest that while afferent proprioceptive feedback of hand position around the target most likely plays a role in adaptation, it is not necessary to induce adaptation.

## Introduction

Imagine a tennis player competing in a long match. As fatigue sets in, the player gradually and unconsciously adjusts their swing to overcome the effects of muscle fatigue and keep their shot toward the intended goal. This ability to unconsciously adapt well-learned movements to achieve a motor goal is known as implicit motor adaptation (Krakauer et al., 2019). Early research has suggested that implicit motor adaptation during visually guided actions is driven by visual sensory prediction errors (SPE) created by the difference between the expected and actual visual feedback resulting from a motor command (Shadmehr et al., 2010).

This hypothesis has typically been formalized through a state-space model where the amount of adaptation is governed by a learning rate (the proportion of error corrected for on a given trial) as well as a retention factor (the proportion of the state estimate retained from trial-to-trial and captures the natural tendency to return to a baseline state). Based on this model, adaptation eventually reaches a state where the response to the visual error (e.g., the difference between the reach goal and visual feedback) and decay of the adapted response reach an equilibrium.

Although a visuocentric state space model has successfully predicted adaptation behaviour during many different paradigms (Huang et al. 2011; Smith et al., 2006, Vaswani et al., 2015), it does not recognize other potential error signals driving adaptation, such as those arising from proprioception.

To address the limitations of a visuocentric view, the Proprioceptive Re-alignment Model (PReMo) has been proposed as an alternative framework for understanding implicit adaptation (Tsay et al., 2022). PReMo frames adaptation as being driven by a proprioceptive error, rather than a visual one, and can successfully account for adaptation phenomena across visuomotor rotation and force field adaptation studies. The model posits that during a goal-directed action, an actor’s perceived hand position results form a combination of vision, proprioception, and the motor goal. When the perceived hand position does not align with the task goal, a proprioceptive error signal is generated. Adaptation then persists until the perceived hand position is realigned with the goal and the error is nullified.

According to PReMo, the perceived hand position is determined mainly by afferent proprioceptive feedback, the proprioceptive goal, and visual input from the goal-directed action. However, in a recent study examining implicit adaptation during target-directed reaching movements in rare individuals with deafferentation (severe, or complete, proprioceptive loss), Tsay et al. (2023) found similar adaptation between deafferented individuals and healthy controls. Previous studies utilizing visuomotor rotations or forcefield perturbations have also shown comparable levels of adaptation between individuals with deafferentation and healthy control participants (Bernier et al., 2006; Sarlegna et al., 2010). These findings suggest that afferent proprioceptive feedback may not be necessary for adaptation, indicating the need to consider other potential information sources.

Indeed, during a reaching movement, the position of the hand can be estimated from a combination of both afferent and efferent sources, such as the prediction of sensory consequences for the given movement derived from the efference copy (Gandevia et al., 2006; Miall et al., 2007; Proske & Gandevia, 2018). The motor-related efferent prediction has been suggested as a mechanism for the error signal driving adaptation in deafferented individuals, as there may be a mismatch between the actual movement outcome and the expected outcome based on the efference copy (Bernier et al., 2006). This highlights the importance of considering a role for efferent signals in implicit adaptation. Given that both afferent and efferent information would be closely aligned during a typical reaching paradigm, current protocols do not allow us to isolate the efferent component and determine its relative contribution towards adaptation. Thus, the purpose of the current study was to examine the effects of dissociating the efferent from the afferent contributions to implicit sensorimotor adaptation.

Here, we utilized a variation of the clamped visual feedback reaching task to address whether implicit adaptation persists when movement outcome information is based primarily on motor-related efferent information. In this task, cursor feedback is “clamped” to a specific angle relative to the target. Participants are fully informed of the perturbation and instructed to aim directly for the target. This procedure has been shown to effectively isolate implicit learning and minimizes any potential contributions from strategic (explicit) processes to overall behavioral change. Critically, instead of participants performing actual reaching movements as they would in the typical paradigm, they produced an isometric force onto a stationary handle to move a cursor towards targets on the visual display. During perturbation trials, the cursor feedback was rotated relative to the target, independent of the actual force movement direction. By applying force in different directions, participants were able to “reach” towards different targets without actually moving their hand. Although afferent proprioceptive feedback of the hand remained at the home location, there is presumably a sensory prediction based on the isometric action which could contribute to perceived hand position with respect to the goal. From this perspective, the efferent-based percept of the generated movement will be influenced by the misaligned visual information, inducing a shift which then drives adaptation (e.g., Rossi et al., 2021). Thus, we predicted that participants would exhibit implicit adaptation during the isometric aiming task by slowly drifting the direction of their force exertion away from the target when exposed to clamped visual feedback. Further, we also predicted that adaptation would reach similar asymptotic levels compared to those found in previous clamped visual feedback reaching studies (e.g., Kim et al., 2018; Morehead et al., 2017; Tsay et al., 2023), based on the notion that the efferent sensory component would compensate for the missing afferent contributions.

## Results

Figure 1 illustrates the experimental set-up and basic protocol. Participants applied force to a stationary handle to move a cursor from a central home location towards a visual target. Participants completed no-feedback and veridical feedback baseline trials, followed by clamped-feedback trials where the cursor trajectory was rotated 15°, and finally no-feedback trials to test for aftereffects. The main dependent measure used to index adaptation was mean force vector angle, calculated as the average angular difference between the target and force-transformed cursor endpoint over all four trials in one cycle (one to each target).

**Figure 1.**
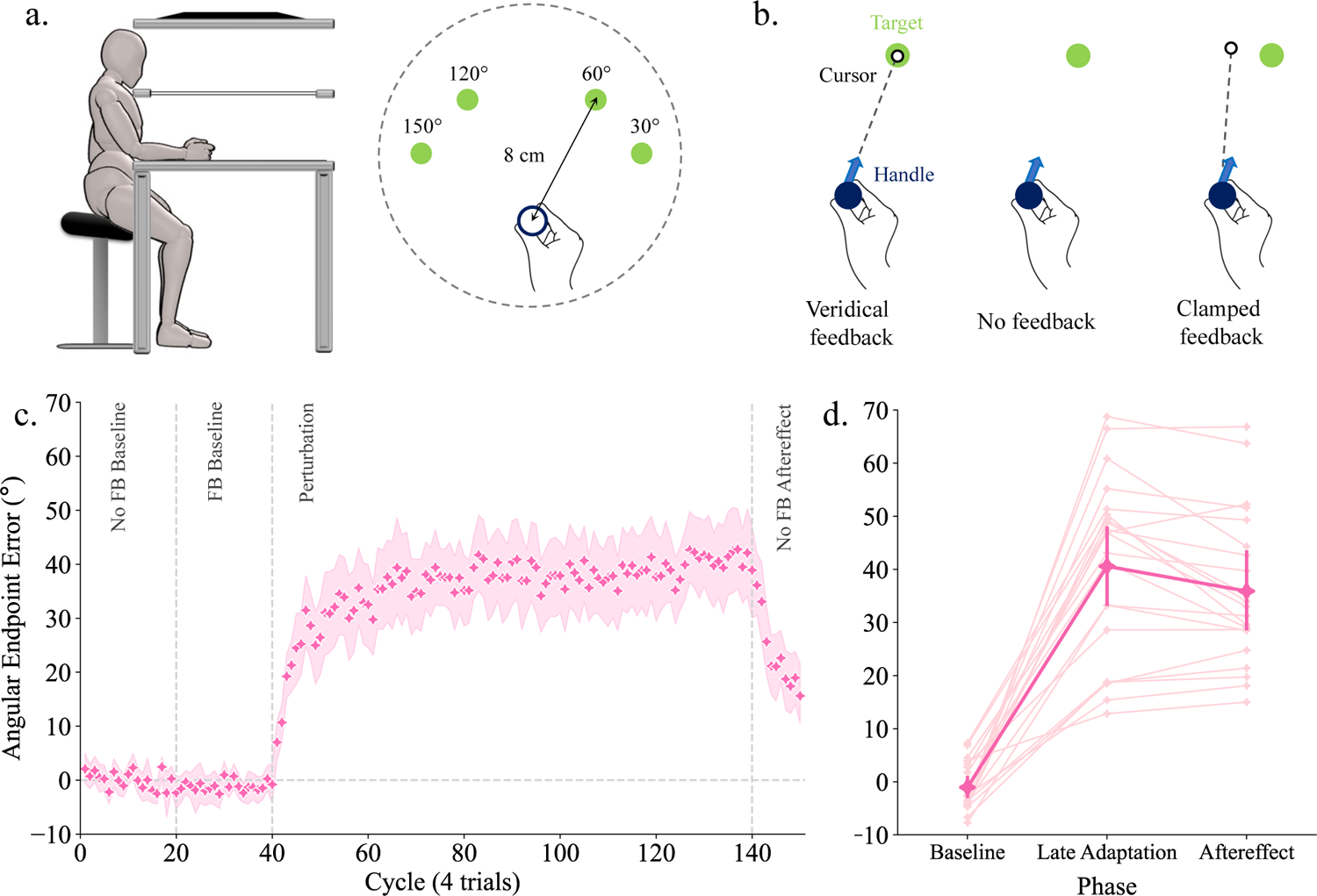
**A**. Experimental Set-up. Participants were seated in a comfortable position while holding the isometric apparatus located on a table surface. Visual feedback was reflected onto a one-way mirror which occluded vision of the actual hand position. Participants exerted force onto the isometric handle to move a cursor on the screen to one of the four possible target locations. The force was translated into cursor motion, such that the direction and speed of cursor movement reflected the force exerted onto the handle. During baseline, participants completed 80 trials with both veridical feedback and no feedback. During exposure, participants completed 400 trials with clamped-feedback where the cursor trajectory was rotated 15°. The final block consisted of 40 no-feedback trials to test for any adaptation aftereffects. **B**. Visual feedback (FB) types. On veridical feedback trials, the cursor moved in the direction of the isometric force exertion. On no-feedback trials, there was no cursor displayed on the screen. On clamped-feedback trials, the cursor was rotated 15° relative to the target, regardless of force direction. **C**. Mean implicit adaptation behaviour during an isometric task (n = 21). Data points represent angular endpoint error averaged across cycles for all participants across all experimental phases. Vertical dotted lines indicate transitions between blocks. Shaded area = 95% confidence interval based on bootstrapped distribution. **D**. Mean angular endpoint error during baseline, late adaptation, and aftereffect phases. Points represent average angular error for each individual participant during each experimental phase. Error bars represent multi-level bootstrapped within-subject 95% confidence intervals. Data plots were generated using Seaborn.

### Implicit adaptation persists in isometric movements

Consistent with our hypothesis, we observed robust adaptation during our isometric task (Figure 1C), with all 21 participants generating progressively larger forces in the opposite direction of the clamped cursor over the course of the perturbation block. Early adaptation occurred at a rate of ∼3.6° [2.4, 4.9] per cycle, with the final magnitude of adaptation eventually plateauing at a mean force vector angle of 40.8° [33.1, 48.4] (Figure 1C). As seen in Figure 1D, there was a clear effect of phase on mean angular error [F(2,40) = 116.225, p < .001, η*_p_*^2^ = .853]. Post-hoc comparisons revealed a marked difference between force angle at baseline versus late adaptation (*p* < .001, *d* = −3.208), as well as the first cycle of the post-test (*p* < .001, *d* = −2.852), which typically serves as the standard measure of implicit learning (Figure 1D). Importantly, there was no reliable difference between late adaptation and the aftereffect (*p* = 0.131, *d* = 0.356). The lack of change in force vector angle when the visual feedback was removed provides additional evidence that the observed adaptation was implicit, and suggests an updating of the forward model during the task.

### Reaction times and movement times remain stable across experimental blocks

Although there was no restriction placed on reaction time through task instruction, participants initiated their movement within 370 ms [334.6, 405.4] on average. These relatively short reaction times lend further credence to the interpretation that the behavioral changes we observed were implicit in nature, as explicit strategies require deliberation and are typically associated with longer reaction times (Fernandez-Ruiz et al., 2011; Haith et al., 2015). A one-way rm-ANOVA comparing average reaction times across experimental phases (baseline no-feedback, baseline veridical feedback, early adaptation, late adaptation, aftereffect) confirmed there was no effect of phase on reaction time [*F*(4,80) = 1.588, *p* = .215, η*_p_*^2^ = .074].

Median movement times averaged across participants over the course of the experiment were 167.3 ms [152.9, 181.8], indicating relatively rapid execution of the force movement. There was an effect of phase on movement time [*F*(4,80) = 14.161, *p* < .001, η*_p_*^2^ = .415]. Movement times during the veridical feedback baseline trials were slightly slower (*x^-^* = 190 ms) than all other experimental blocks (*p* < .001), likely due to the effect of veridical visual feedback on movement control. There were no other reliable differences between movement times among the other experimental blocks.

### Enhanced adaptation supports a mechanism beyond vision

Our results demonstrated that implicit adaptation is preserved in a goal-directed isometric aiming task. When the perturbed visual feedback was introduced, there was a rapid shift of force direction in the opposite direction of the clamped cursor until a plateau was reached. Moreover, when the perturbed feedback was removed, participants continued to initially direct their force exertion away from the target, showing implicit aftereffects. These findings are consistent with our proposal that, even without dynamic reach-relevant proprioceptive feedback, an efferent-based sensory prediction of the action can still contribute to a perceived action outcome. In our view, this efferent-based estimate, and the presumed bias induced by the misaligned visual feedback, formed an error signal to drive adaptation. These results support our hypothesis, providing evidence for the potential contribution of efferent information towards implicit adaptation.

Our interpretation of the results remain predicated on the premise of a proprioceptive realignment mechanism, such as that proposed by PReMo, with the inclusion of an efferent-based prediction that contributes to the estimate of the action outcome. However, the results from this experiment do not necessarily rule out a visuocentric interpretation. The task constraint of the hand remaining at its central location minimized the contribution of proprioceptive feedback in providing information of hand position with respect to the target, as was our intention. But this might also have led to the dominance of visual feedback (e.g., Berniker & Kording, 2011). Thus, the persistence of the visual error might have been expected to induce adaptation similar to a typical reaching paradigm. However, our results also demonstrated enhanced adaptation in an isometric task compared to previous findings using a clamped-feedback reaching paradigm, suggesting that vision alone may not explain the observed behaviour.

Although previous studies using reaching tasks usually report an asymptote of adaptation around 20°, they typically use a full array of targets spread throughout the entire 360° workspace around the home position (H. E. Kim et al., 2018; Morehead et al., 2017; Tsay et al., 2020). To confirm that the heightened adaptation in our task was not simply a result of our target display, and to be able to draw further comparisons to typical reach adaptation, we tested an additional twelve participants in a follow-up experiment. All experimental parameters were the same as the main experiment, except participants performed actual reaching movements instead of isometric force exertions. Participants were instructed to shoot through the target before returning to the home position to begin the next trial. This allowed us to directly compare the adaptation between an isometric paradigm and a reaching paradigm. Critically, this also provided a test of a visuocentric account, which would predict comparable adaptation between the two task types, as the magnitude of visual error was kept constant across the isometric and reach tasks.

### Reaching adaptation was reduced compared to isometric behaviour

Similar to previous research, robust adaptation was observed when participants reached towards a target with clamped visual feedback (Figure 2A). All participants demonstrated a clear deviation of force vector angle away from the perturbed feedback, eventually reaching a plateau. Early adaptation occurred at a rate of 1.7° [0.5, 2.9] per cycle, and plateaued at around 11.9° [8.7, 15.3]. Although this asymptote of adaptation occurred at a smaller angle than previous studies, it confirmed that the enhanced adaptation observed in the isometric task was not simply due to the target set-up and protocol. Angular endpoint error remained similar on the first cycle of the post-test at approximately 11.3° [8.35, 14.17], indicating the presence of an aftereffect.

**Figure 2.**
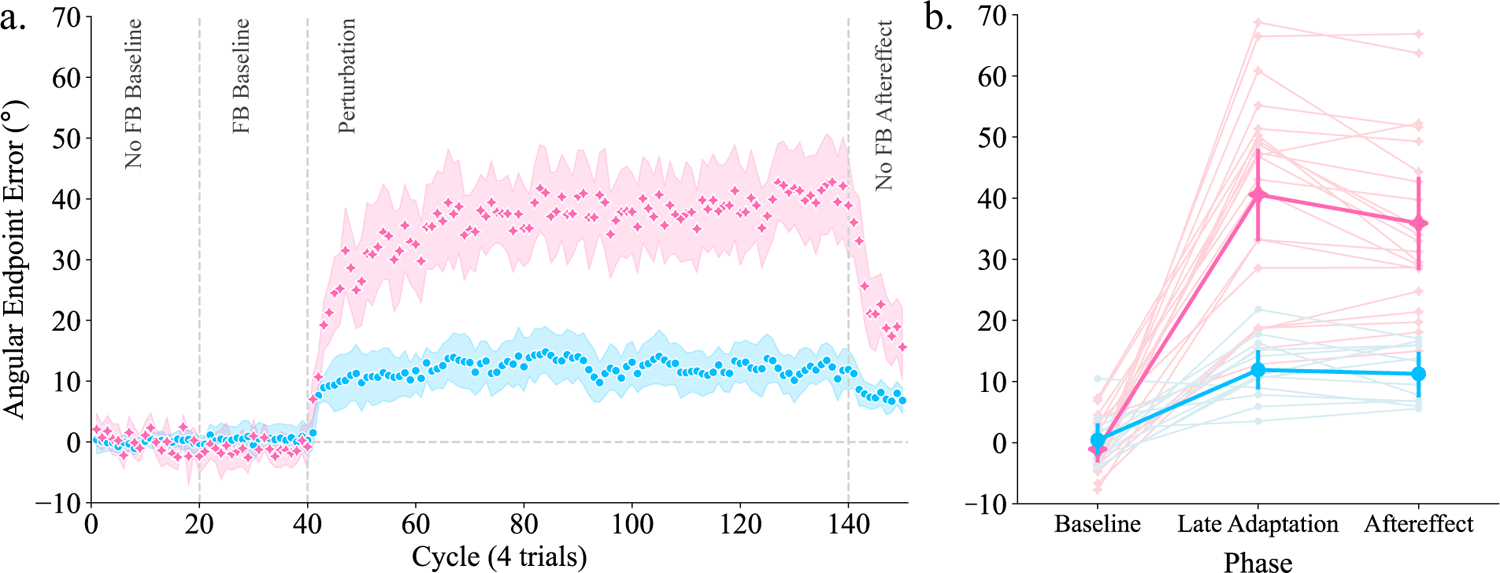
**A**. Implicit adaptation differs between isometric and reaching tasks. Average endpoint error over cycles for all participants in the isometric (pink stars) and reaching (blue circles) group. Shaded areas = 95% confidence intervals based on bootstrapped distribution. **B**. Angular endpoint error during baseline, adaptation, and aftereffect measures. Pink = isometric group, blue = reaching group. Points represent each individual participant. Error bars represent multi-level within-subject bootstrapped 95% confidence intervals. Data plots were generated using Seaborn.

As seen in figure 2A, there was an effect of phase (baseline, adaptation, aftereffect) on mean angular error [*F*(2,22) = 23.292, *p* < .001, η^2^_p_ = .679]. Post-hoc comparisons showed a clear difference between angular error in the baseline phase compared to the end of adaptation (*p* < .001, d = −2.468) as well as the first trial of the aftereffect phase (*p* < .001, d = −2.311). There was no reliable difference between angular error during adaptation compared to aftereffect (*p* = .701, d = 0.157), further confirming that the adaptation observed was implicit.

To confirm the difference in adaptation between the isometric group and the reaching group, we submitted angular error to a group (isometric or reaching) vs phase (baseline, adaptation, aftereffect) mixed-design ANOVA. There was a significant Group * Condition interaction [*F*(2,62) = 22.709, *p* < .001, η*_p_*^2^ = .493]. While both groups performed similarly during baseline (*p* = 1.000, d = −0.138), there was a marked difference between groups during both the adaptation (*p* < .001, d = 2.657) and aftereffect conditions (*p* < .001, d = 2.297), demonstrating that the isometric group had greater adaptation and aftereffects compared to the reaching group (Figure 2B).

Experiment 2 demonstrated that during dynamic reaching, our task protocols did not lead to comparable levels of adaptation as observed in Experiment 1, further supporting our finding that implicit adaptation was significantly heightened in a goal-directed isometric task. These results confirm that the increased adaptation in Experiment 1 was not simply due to the specific target placements used. More importantly, the findings do not support the visuocentric view of implicit adaptation, as different amounts of adaptation were induced in two tasks which shared identical visual stimuli.

## Discussion

The purpose of this study was to examine the effect of dissociating the afferent and efferent contributions to implicit sensorimotor adaptation. We utilized a unique isometric aiming task where participants exerted force onto a static handle to advance a cursor towards targets on a visual display. By making force movements without actually moving the limb, we were able to examine the potential contribution of the movement-related efferent prediction without the confounding influence of proprioceptive feedback tied to the action (cf. O. A. Kim et al., 2022; Mostafa et al., 2019). Although we did not directly remove proprioceptive feedback, we were able to dissociate the feedback of hand location from the efferent prediction of a typical reaching movement.

We found that participants implicitly adapted in a task where they were required to generate isometric forces to move a clamped cursor on a visual display instead of performing actual reaches to the target. Furthermore, we observed greater overall adaptation than in a typical clamped reaching paradigm (H. E. Kim et al., 2018, 2019; Morehead et al., 2017). This finding was confirmed in our secondary experiment where participants performed actual reaching movements and demonstrated significantly less adaptation. These findings suggest that while proprioceptive feedback of hand position at the target location most likely plays a role in adaptation behavior, it may not be necessary in and of itself to induce adaptation.

Our original question for this study was motivated by PReMo, which advanced the idea that proprioception is a key component of implicit adaptation. More specifically, we wanted to examine what information sources contribute to the perceived hand position and adaptation in a target-directed movement. PReMo postulates that the visually recalibrated estimate of hand position is based on proprioceptive feedback and gives rise to the “proprioceptive shift”, the perceived misalignment between the hand position and target, that drives implicit adaptation. Thus, would the absence of proprioceptive feedback, or its dissociation from the action as in our isometric task, preclude a proprioceptive shift and therefore negate implicit adaptation? Our results clearly show this not to be the case, as adaptation prevailed in the absence of reaching.

Alternatively, we might have expected unbounded adaptation in the isometric task based on PReMo. While the perceived hand position is composed of proprioceptive feedback, vision, and the motor goal during a typical reaching task, it may only be composed of the latter two during the isometric task, resulting in unbounded adaptation to offset a constant shift towards the visual perturbation. Considering adaptation reached a clear plateau during the isometric task, we must consider other potential sensory signals, such as the efferent motor prediction, which may be contributing to implicit adaptation behaviour.

Along these lines, Zhang et al. (2024) have recently proposed the Perceptual Error Adaptation (PEA) model that also incorporates perceived hand position as a key feature in the mechanism to explain implicit adaptation. The model proposes that when a participant reaches towards a target with error-clamped feedback, a perceptual estimate of hand position is formed from the combination of three sensory cues: visual feedback of the cursor, proprioceptive feedback from the actual hand position, and the motor prediction of the command towards the target, with each cue weighted based on its relative uncertainty. The discrepancy between the estimate of hand location and the target gives rise to a perceptual error that drives subsequent adaptation.

In our isometric task, even though proprioceptive feedback is not actually removed, it no longer contributes relevant information to the perceived hand position with respect to the target since the hand is not moving. Consequently, the relative uncertainty of proprioceptive input is presumably much higher. From the perspective of the PEA model’s cue combination mechanism, this would result in a larger relative weighting of the visual cursor, thereby causing a greater perceptual shift towards the cursor and driving increased adaptation compared to reaching.

Although this prediction aligns with our experimental findings, it assumes that dissociating proprioceptive information simply results in increased proprioceptive variability. Furthermore, similar to PReMo, this model neglects to recognize the potential influence of the sensory prediction based on the actual outgoing motor command.

Based on the findings from the current study, we suggest that models of implicit motor adaptation should include motor-related efferent contributions (Figure 3). While we acknowledge that both the PReMo and PEA models do posit a form of a motor-related prediction based on the motor command, it is distinct from what we are proposing. The motor-related prediction from these models is always associated with the intended aim towards the target – i.e., the motor goal. By this definition, the motor prediction acts like a prior, and is associated with an assumed outcome at the goal. While this may be the explicit sensory prediction as the task instruction is to continue reaching towards the target despite perturbed visual feedback, we argue that there is also an efferent sensory prediction based on the actual motor command that is actually generated and executed. This is separate from a higher-level expectation of the movement outcome based on the intended aim. When the end result of an executed reaching movement differs from the explicitly intended goal (e.g., during late adaptation), the efference-copy based prediction would be with respect to where the hand actually went and can therefore contribute to the perceived hand position.

**Figure 3.**
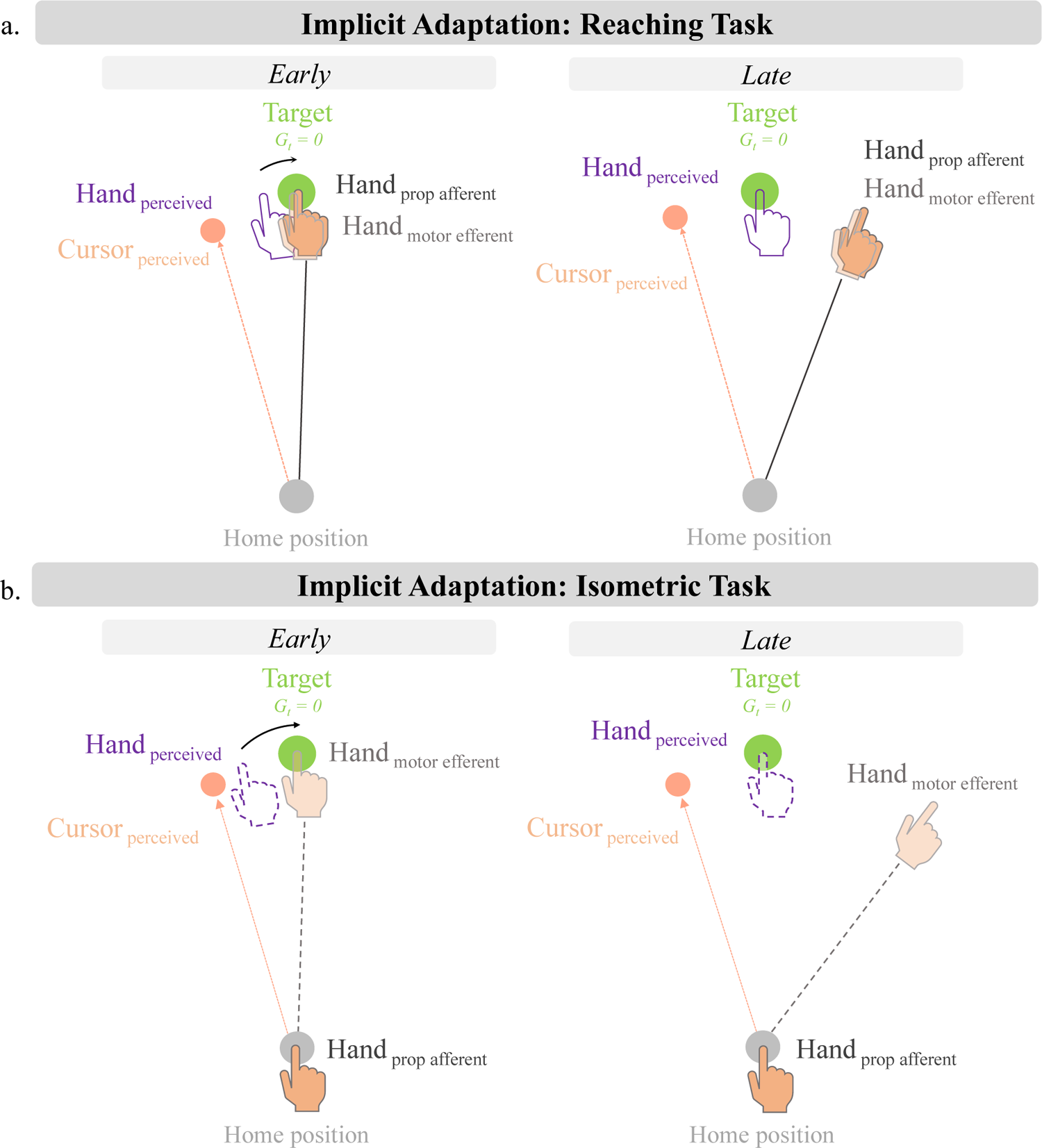
**A.** Implicit adaptation during a clamped feedback reaching task. We propose that the perceived hand position is derived from vision, the motor goal, afferent proptioceptive information, and efferent information tied to the outgoing motor command. The perceived hand position is initially biased towards the visual cursor which drives adaptation of reach direction. **B.** Implicit adaptation during an isometric task. While afferent proprioceptive information is no longer contributing to perceived hand position, there is still the efferent-based motor prediction which can contribute, and therefore play a role in adaptation. The discrepancy between the perceived “hand” position and the target drives adaptation of the force trajectory away from the clamped cursor.

Considering the efferent-related motor prediction contributes to position sense, we suggest that this information source be considered towards the proprioceptive contribution of the perceived hand position. Currently, in both the PReMo and PEA models, the proprioceptive estimate of hand position is based on afferent information only. However, studies have shown that our estimate of hand position is also based on the sensory prediction derived from the efference copy (Gandevia et al., 2006; for review see Proske & Gandevia, 2018). Thus, we propose that these models should incorporate both the afferent and efferent components of proprioception when considering proprioceptive contributions.

Within the framework of PReMo, the proprioceptive estimate of the hand would now be composed of two components: the afferent proprioceptive input and the efferent-based sensory prediction based on the executed motor command (Figure 3A). In the isometric task, while the afferent input may no longer directly contribute to position sense with respect to the target, the efferent-based sensory prediction would still contribute to the perceived movement outcome (Figure 3B). This efferent-based percept of the hand position (combined with the motor goal towards the target) would then be biased by the visual cursor to create the error signal which would drive adaptation. The addition of the efferent contribution provides a cue of hand position or reach trajectory even in the absence of proprioceptive feedback, allowing for a “proprioceptive” (or perceptual) shift which could drive subsequent adaptation (see also Rossi et al., 2021). Since the proprioceptive estimate of the hand would be based solely on the efferent component during the isometric task, there could be a heavier influence of vision when determining the perceived movement outcome, based on ideas of optimal cue integration as in the PEA model, leading to an increased error signal driving heightened adaptation.

Our current work and proposed revision to include an efferent-based contribution supports Tsay et al.’s (2024) recently updated formulation of PReMo and its corresponding predictions for adaptation behaviour. Moreover, our prediction aligns with those made by these authors when testing individuals with proprioceptive loss in the clamped visual feedback task. Similar to the isometric task, these individuals would not have an afferent contribution towards their estimate of hand position, but presumably must rely on a motor prediction to estimate the outcome of their movements. The findings showing that adaptation reached a clear plateau in the individuals with proprioceptive loss (as opposed to following an unbounded trajectory), supports the notion that the efferent motor signals can play a role in implicit motor adaptation.

With respect to the PEA model, the incorporation of the efferent contribution in a manner similar to what we proposed for the PReMo model, would provide a theoretical basis for increased proprioceptive uncertainty when afferent input is not available, or made task-irrelevant. In our isometric task, only the efferent component would contribute to position sense. Thus, compared to a situation where both afferent and efferent sources contribute to the proprioceptive estimate of hand location, the absence (or attenuation) of the afferent contribution (or conversely, the absence of the efferent contribution), should result in a less reliable proprioceptive estimate. This would result in a greater weighting of the visual input and a greater perceptual shift towards the visual cue, leading to heightened adaptation (Figure 3B). Thus, the addition of the efferent-based proprioceptive information provides a theoretical basis for the increased proprioceptive variability in the isometric task and subsequent increased adaptation.

Our proposal of an efferent contribution is not only consistent with what is known about the proprioceptive, or kinesthetic, senses (Proske & Gandevia, 2012, 2018) but also with previous work that have postulated an efferent-based contribution to hand localization and adaptation behaviour (Henriques & Cressman, 2012; Rossi et al., 2021). Previous studies have isolated afferent feedback-based, contributions from motor-related efferent contributions by comparing robot-driven passive movements to active movements (Mostafa et al., 2019; ‘t Hart & Henriques, 2016). These studies have shown that while implicit reaching aftereffects can be observed after both active and passive training to a visuomotor rotation, a larger proprioceptive shift is induced after active training, suggesting that the updated efferent prediction contributes to hand localization. In addition, Bernier et al. (2006), in a study of a proprioceptively deafferented individual adapting to a visuomotor rotation, has also suggested that adaptation is driven by a sensory prediction error created between the actual consequences of a movement, and the predicted consequences based on the outgoing motor command. The results from these studies, as well as the current findings, suggest that the motor-related efferent prediction plays an important role in implicit motor adaptation, and should therefore be incorporated into current computational models.

The purpose of this study was to examine the contributions of efferent compared to afferent signals towards implicit sensorimotor adaptation. To the best of our knowledge, this was the first study to isolate implicit adaptation during an isometric task. The results offer insights into the potential sensory cues needed to induce adaptation and allow us to critically examine the current models of implicit adaptation. While these findings are novel and important towards our greater understanding of implicit adaptation, various future studies are still needed to confirm the driving mechanisms for adaptation. Our data cannot directly reveal if any perceptual realignment occurred during the course of adaptation, and if so, whether or not the perceptual error was fully nullified. Studies examining these questions will allow for further refinement of models of implicit adaptation and expand our understanding of sensorimotor adaptation more broadly.

## Methods

### Experiment 1

#### Participants

Thirty-three students (9 male, 24 female; ages 18-27) from the University of British Columbia were recruited to participate in this study. Twenty-one individuals participated in the main experiment while the remaining twelve completed a follow-up study. Participants were free of neurological or musculoskeletal injury preventing them from performing movements with their dominant limb, were naïve to sensorimotor research, and had normal or corrected-to-normal vision. All participants were asked to provide informed consent prior to participation and were compensated financially for their time. All study protocols were approved by the UBC Behavioural Research Ethics Board.

#### Experimental Setup

Participants performed the isometric reaching task while seated in a comfortable position. Targets were displayed on a 27-inch LCD monitor (600 x 339 mm display, 240 Hz refresh rate, 1920 x 1080-pixel resolution) and reflected onto a one-way mirror located under the screen to provide an illusion of the targets being located on the table surface. The position of the mirror and dark environment of the experimental room occluded peripheral vision of the arm.

The isometric handle used to carry out the experimental task was placed approximately 36cm from the edge of the table surface below the mirror. The handle used was custom built using a small sized 3-axis force sensor (USL06-H5-50N-E; Tec Gihan). The sensor was 20×20mm and had a range of 25N in the Fx and Fy direction (although the sensor also detects force in the Fz direction (vertical), we were less concerned with this direction as the participant exerted forces in the horizontal plane). A custom 3D-printed handle (16-sided, 2.3 cm diameter, 5 cm height) was mounted onto the sensor such that sheer force applied to the handle was encoded by the sensor to determine force direction.

The forces exerted onto the handle were ultimately transformed and displayed on the screen as cursor motion. When a force was applied to the handle, analog signals from the force sensor were passed through an instrumentation amplifier (EI-1040; Electronic Innovations Corp.) at 1000x gain. Next, analog output signals from the instrumentation amplifier were passed through an analog amplifier at 10x gain and bandpass filtered DC to 10Hz. This amplifier also allowed the experimenter to manually adjust the voltage offset to deal with any drift in the sensor. This amplified and filtered voltage signal was then recorded through a 16-bit analog-to-digital unit of an OPTOTRAK (ODAU II, Northern Digital Inc) 3D motion capture system sampling at 500 Hz. This signal was transformed using the force sensor’s calibration matrix to return the force values for Fx, Fy, and Fz in Newtons. This was done through software written in MATLAB (R2015b, The MathWorks Inc). The force values were then tranformed such that 1N translated to ∼12mm (approximately 36 pixels; 1mm ∼ 3 pixels) of cursor movement on the screen. The ODAU II was polled continuously (providing an effective sampling rate of ∼250Hz) to provide the visual cursor feedback.

#### Experimental Task

Participants were asked to apply force to the handle to move a cursor towards targets (green circles; 6 mm diameter) presented on the visual display. The cursor (white circle; 3.5 mm diameter) always began in the home position, indicated by an empty circle (10 mm diameter).

The home circle appeared yellow while the cursor was inside, and red once it left. After a given 500ms foreperiod, a target was presented in one of four locations in front of the home position (30°, 60°, 120°, 150°) at a radial distance of 8cm. Participants were instructed to prioritize both speed and accuracy when completing the task. Movement time, defined from the time the cursor left the home circle until it crossed the target perimeter, was required to be between 100-400ms. This short movement time limited the participants from making online corrections during the movement. Movement times outside of this window triggered an auditory tone signalling an error, and the experimenter provided verbal feedback on whether the movement was too slow or too fast. After the movement was complete, participants relaxed, and the cursor reappeared in the home circle to begin the next trial.

#### Feedback Types

On veridical feedback trials, cursor feedback was given to the participant such that the cursor direction was directly coupled with the direction of force applied to the handle. On no-feedback trials, participants did not see a cursor while directing force efforts towards targets. On clamped feedback trials, cursor feedback was rotated 15° clockwise or counterclockwise (counterbalanced between participants) relative to the target. Thus, no matter what direction the participant applied force to the handle, the cursor always followed the same path. It is worth noting that the amplitude and speed of the cursor movement was still tied to the participant’s force, such that a larger force resulted in a faster-moving cursor. Since the direction of the cursor was uncoupled to the participant’s force exertion, they were instructed to ignore this cursor feedback and continue generating forces that were directed towards the target. On trials where visual feedback was present, the cursor remained visible for 200 ms after the participant crossed the target perimeter.

For the first 35 mm of the radial distance from home to the target, a position-based mapping was used for this portion of the cursor display, where the forces recorded were directly mapped to the position and movement of the cursor (e.g., 1N ∼ 12 mm). For the last two thirds of the radial distance, the cursor entered an “automatic zone.” The last 5 samples collected from the non-automatic zone were averaged and used to calculate the angular direction of the resultant force, which determined the cursor trajectory for the remainder of the cursor movement out towards the target. This calculation was done online such that the cursor appeared to seamlessly move from one zone to the next. The automatic zone helped to ensure that the cursor movement was smooth out toward the target, and that online corrections were mitigated. In other words, whatever force direction was exerted during the first third of the movement determined the direction of the remainder of the movement, regardless of a change in force direction during the latter part of the trial. Therefore, even if online corrections did occur despite the fast movement time, they did not manifest through the cursor display. The speed of the cursor movement however was still tied to the force exertion such that a stronger push moved the cursor in a faster manner. This cursor mapping also helped reduce the exertion effort needed from the participant. In attempt to make the pushing movement as similar to a regular reach as possible, the participant did not have to over-exert themselves when pushing on the handle, just as they would not have to use much force to move their hand to a target in a typical reaching paradigm. This cursor mapping addressed this issue by seamlessly advancing the cursor with a relatively small force (∼4N) applied to the handle.

#### Experimental Blocks and Conditions

The experiment consisted of 12 blocks. Each block contained a designated number of cycles. One cycle was defined as four “reaching” movements: one to each target position presented in a randomized order. Participants had the opportunity to explore the nature of the experimental apparatus in a calibration block prior to testing. This block was 60 seconds long, allowing the participant to become familiar with the required force movement as well as the cursor sensitivity. A home position and an arc were drawn on the screen for the entire block. The arc was a white semicircle covering the upper-right and upper-left quadrants and had a radius of 10 cm from home. The participant was instructed to move the cursor in any direction they chose around the arc. They were encouraged to move in various directions so that they could explore the entire workspace. The calibration block was run more than once if the participant felt they needed more time to familiarize themselves with the apparatus. Once comfortable with the task, they participated in a practice block of 60 targeted reach trials with veridical feedback. The first 40 practice trials were divided into 10 trials per target, where all 10 trials to each target were presented sequentially. This gave the participant an opportunity to repetitively aim to the same location and develop familiarity with exerting force towards the given target. The remaining 20 practice trials were pseudorandomized such that each target was presented once before they were repeated. Practice blocks were repeated if the standard deviation of their cursor angle over the whole block was greater than 20 degrees (indicating large variability in performance), or they did not feel comfortable with the task. Once they confirmed they could confidently aim to the different targets, the experimental trials commenced.

Participants began the experimental trials with two baseline blocks. Baseline block 1 consisted of 80 no-feedback trials, and baseline block 2 consisted of 80 veridical feedback trials. These blocks served as an assessment of individual baseline bias and variability.

Next, participants performed a practice block of clamped feedback to familiarize themselves with the nature of the cursor movement. On the first trial, participants were shown a target straight in front of them and were asked to execute a force exertion towards that target.

After seeing the rotated cursor feedback, the experimenter explained that the cursor feedback would remain rotated relative to the target for the next series of trials. To demonstrate this to the participant, they were presented with the same target on the next trial but were instructed to exert a force in the rightward direction. Next, they were asked to exert a force in the leftward direction. On the final practice trial, they were asked to exert a force in any direction of their choosing.

After observing the same cursor feedback on all four practice trials no matter what force was exerted on the handle, the experimenter confirmed with the participant that they understood the nature of the cursor feedback before advancing to the exposure blocks.

The next five blocks (80 trials each) consisted of clamped cursor feedback. Participants were instructed to ignore the feedback and continue making force exertions towards the target. Participants were given the opportunity for a short break between blocks to prevent mental fatigue. The final block consisted of 40 no-feedback trials towards the targets to assess aftereffects. The participants were told to keep aiming towards the targets as in the exposure blocks, but there would no longer be any feedback displayed on the screen.

#### Data Collection and Measures

The main dependent measure in this experiment was the mean force vector angle, which was used to index whether there was adaptation. Mean force vector angle was calculated as the average force vector angle across all four trials in one cycle. Force vector angle on each trial was computed as the angular difference between the target and the cursor endpoint. In order to identify outliers, a 5-point moving average was calculated for each participant and target location. Trials were removed from analysis if they were greater than 2.5 standard deviations from the moving average, which resulted in 1.92% of trials being removed. An additional 2.83% of trials were removed due to movement time violations or other trial errors. Average angular error during baseline trials for each individual participant was subtracted from all trials to account for inherent biases in aiming angle. Early adaptation rate was measured as the average force angle from the first five cycles of clamped feedback trials. Late adaptation was measured as the average angular error over the last 10 cycles of clamped feedback. The aftereffect was measured as the average force angle over the first cycle of no-feedback trials. In addition to mean force angle, baseline force angle variability was calculated as the standard deviation of participants’ force angles during the no-feedback baseline trials.

Other performance measures were also taken to analyze performance. Reaction time was measured as the time between target presentation and when the cursor left the home region. Movement time was taken as the time between when the cursor left home and when it crossed the target perimeter. Although movement time was restricted to 100-400ms, reaction time was not restricted. Participants were simply instructed to push towards the target as soon as it appeared. Median reaction and movement times were taken for each participant for all no-feedback baseline trials, all veridical feedback baseline trials, the first and last 10 cycles of clamped feedback trials (early and late adaptation), and all no-feedback post-test trials.

To determine if adaptation had occurred, a one-way repeated measured analysis of variance (rm-ANOVA) was used to compare angular error between baseline, adaptation, and aftereffect phases. Significant effects were followed up with Holm corrected *t*-tests to determine how angular error differed in each phase. Separate rm-ANOVAs were used to determine if there were any differences in reaction or movement time across experimental phases (baseline no-feedback, baseline veridical feedback, early adaptation, late adaptation, aftereffect). Significant effects were followed up with Holm corrected *t*-tests to determine how reaction and movement time differed in each of these five phases. Statistical significance was set to *p* < .05 for all measurements and a greenhouse-geisser correction was applied if the assumption of sphericity was violated. Uncorrected degrees of freedom are reported in the text.

### Experiment 2

Instead of using the isometric handle, participants made actual slicing movements through targets on the visual display. Participants wore a splint on their right index finger with an infrared marker at the tip. An OPTOTRAK (3020, Northern Digital Inc) 3D motion capture system was used to track reaching movements at a sampling rate of 500Hz. Participants were instructed to move their hand and finger to the home position to begin each trial. Once they placed their finger/cursor within 30 mm of the home position, an elliptical ring appeared to help guide participants toward the home position without indicating the actual finger position. Once their finger was within 20 mm of home, the cursor appeared. Once the finger was at the home position for 500 ms, the target (6 mm diameter) appeared. All other experimental parameters (target location, experimental blocks, trial numbers) were identical to the main experiment.

## Acknowledgments

The authors would like to acknowledge Gregg Eschelmuller and Nick Butler for their useful comments on the manuscript and helpful discussion throughout this work.

